## Supporting information 1 for "Sloppiness: fundamental study, new formalism and its application in model assessment"

Additional Examples

September 2022

### 1 A pharmacokinetc model cholesterol distribution and turnover

In this example we consider a two compartment pharmacoknetic model given in [1]. This model has been widely used to represent the dynamics of distribution and turnover of labeled cholesterol in humans with impulse injections and then concentration is measure in compartment one [2]. The model has two states, four parameters (Fixing the volume of the compartment ($V$) to unity) and one output. The model diagram is given in Fig. 1 and equations are given in (1).

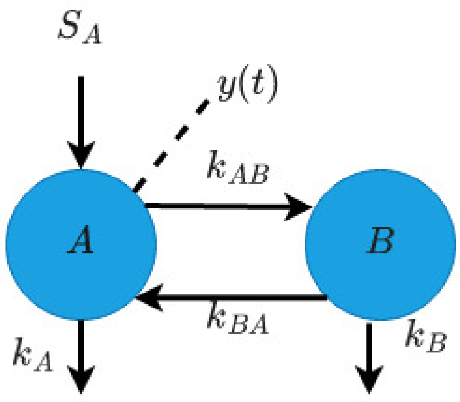


Figure 1: Two compartmental model for cholesterol distribution and turnover

$$\mathcal{M}:\begin{cases}\dot{A}(t) = -(k_A + k_{BA})A(t) + k_{AB}B(t) + S_A(t)\\ \dot{B}(t) = -(k_B + k_{AB})B(t) + k_{BA}A(t)\\ y(t) = \frac{A(t)}{V}\end{cases} \tag{1}$$

The nominal parameter $\theta^* = \begin{bmatrix} 0.5 & 0.25 & 1.1 & 0.75 \end{bmatrix}$. The model output $y(t)$ is simulated for $t = 0$ to $t = 3$ hours with the initial conditions $x_1(0) = x_2(0) = 0$ and the input $u(t)$ is unit impulse input given at $t = 0$.

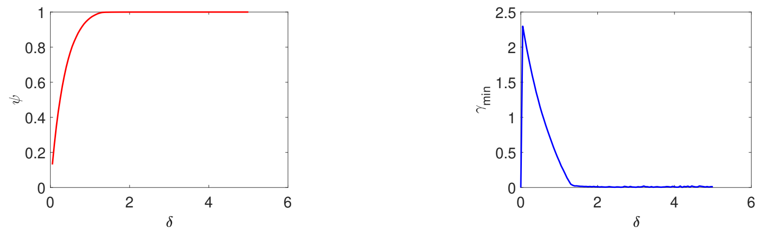


(a) $\delta$ vs $\gamma$ plot

(b) $\delta$ vs $\gamma_{min}$ plot

Figure 2: (a) The curve hits unity value for $\delta > 1.6$ indicating local structural unidentifiability (b) The $\gamma_{min}$ curve also indicate local structural unidentifiability because of zero value of $gamma_{min}$ for non-zero $\delta$

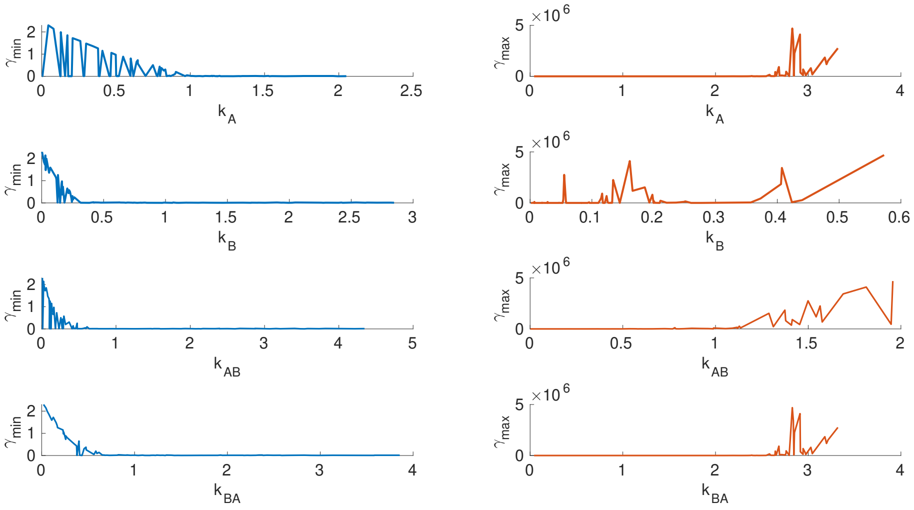


Figure 3: All the parameters have zero value $\gamma_{min}$ and $\gamma_{max}$ for non-zero $\delta_{theta_i}$

From Figs. 2a and 2b, we can clearly see that the model sensitivity curve ($\psi$) hits unity value and $\gamma_m in$ hits zero value for non-zero delta indicating local structural identifibility. Further, to identify which parameters are unidentifiable $\delta_{\theta_i}$ vs $\gamma_{min}$ plot is generated in Fig 3. From Fig. 3 it is evident that all the parameters are unidentifbale for the given model and output pair. This result is consistent with the findings in [1].

### 2 High dimensional biochemical pathway model

A linear biochemical pathway with fourteen states, sixteen parameters and one input is considered for demonstration [3]. The states $x_1(t)$ and $x_{14}(t)$ are measured. The nominal parameter set ($\theta^*$) is taken from [3].

$$\mathcal{M}: \begin{cases} \dot{x}_1 = -\frac{v_m x_1}{k_m + x_1} + p_1 u \\ \dot{x}_2 = -p_1 x_1 - p_2 x_2 \\ \dot{x}_3 = -p_2 x_2 - p_3 x_3 \\ \dot{x}_4 = -p_3 x_3 - p_4 x_4 \\ \dot{x}_5 = -p_4 x_4 - p_5 x_5 \\ \dot{x}_6 = -p_5 x_5 - p_6 x_6 \\ \dot{x}_7 = -p_6 x_6 - p_7 x_7 \\ \dot{x}_8 = -p_7 x_7 - p_8 x_8 \\ \dot{x}_9 = -p_8 x_8 - p_9 x_9 \\ \dot{x}_{10} = -p_9 x_9 - p_{10} x_{10} \\ \dot{x}_{11} = -p_{10} x_{10} - p_{11} x_{11} \\ \dot{x}_{12} = -p_{11} x_{11} - p_{12} x_{12} \\ \dot{x}_{13} = -p_{12} x_{12} - p_{13} x_{13} \\ \dot{x}_{14} = -p_{13} x_{13} - p_{14} x_{14} \\ y(t) = x_1(t) + x_{14}(t) \end{cases} \tag{2}$$

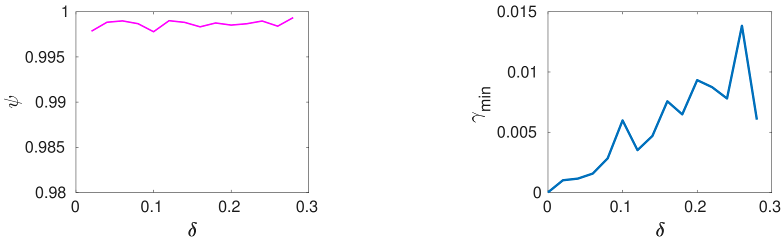


(a) $\delta$ vs $\psi$ plot

(b) $\delta$ vs $\gamma_{min}$ plot

Figure 4: (c) The curve does not hit the unity value indicating local structural identifiability. (b) The curve has significantly small slope and numerically small $\gamma_{min}$ which indicate the sloppiness for the given $\delta$

From Fig. 4b, the model is locally structurally identifiable. In Fig. 4a, the ratio of values of $\frac{\gamma_{max}}{\gamma_{min}}$ increases as $\delta$ increases; the ratio is numerically significant indicating the model is sloppy in the traditional and multi-scale notion of sloppiness. In addition to that, in Fig. 4b, the for $\delta > 0.25$, the $\gamma_{min} \approx 0.015$. This observation implies that the system is $(\epsilon, \delta)$ sloppy, and a sub-set of insensitive parameters will contribute to the sloppiness.

From Figures 5 to 8, it is observed that the parameters $v_m, k_m, p_7$ and $p_9$ are insensitive for $\delta_i > 0.03$ compared to other parameters. The sensitivity in maximum deviation direction is significantly high for the parameters $p_5, p_6, p_{13}$ and $p_{14}$ in the vicinity of $\theta_i^*$. In sum, the biochemical network considered is structurally locally identifiable, and only 4 out of 16 parameters are insensitive for a particular region for the given $\delta$. Further, the model has a moderate multi-scale sloppiness ratio inside the specified $\delta$, which indicates an acceptable isotropic sensitivity in the parameter directions.

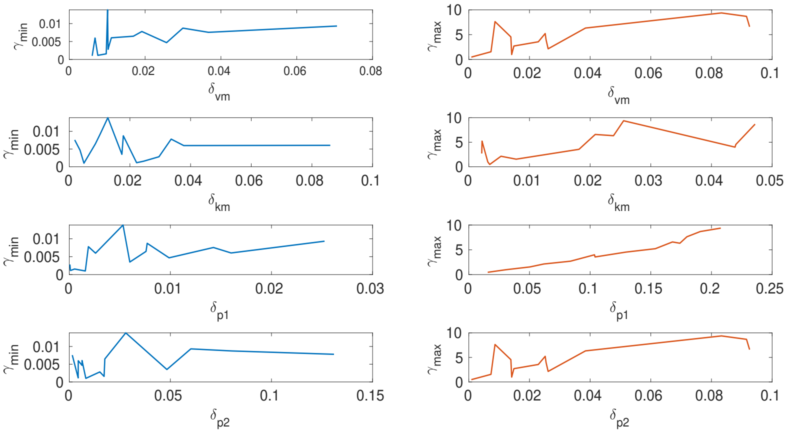


Figure 5: $x$-axis depicts the relative distance of the particular parameter $\theta_i$ from its reference value $\theta_i^*$. $y$-axis on the left is the minimum sum-square deviation and on the it is the right maximum sum-square deviation from $y^*(t)$. Parameters $v_m$ and $k_m$ are insensitive for $\delta_i > 0.03$. The maximum sensitivity of parameter $p_1$ is linearly increasing as $\delta_i$ increases.

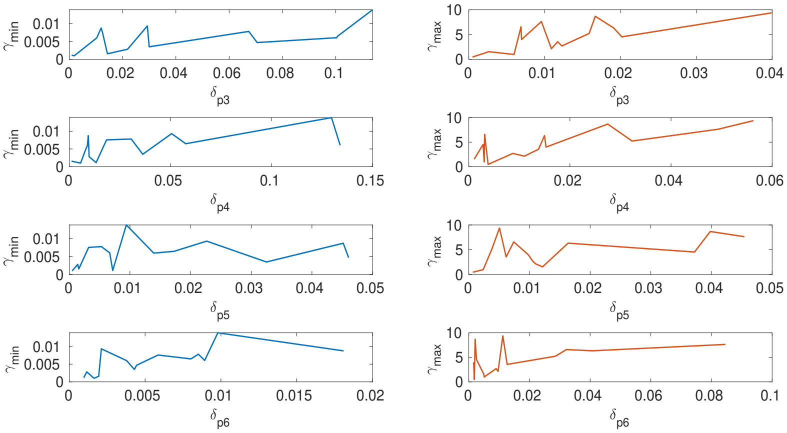


Figure 6: $x$–axis depicts the relative distance of the particular parameter $\theta_i$ from its reference value $\theta_i^*$. $y$–axis on the left is the minimum sum-square deviation and on the right it is the maximum sum-square deviation from $y^*(t)$. While non of the parameters are insensitive, parameters $p_5$ and $p_6$ are highly sensitive in the vicinity of the $\theta_i^*(\delta_i^* < 0.01)$

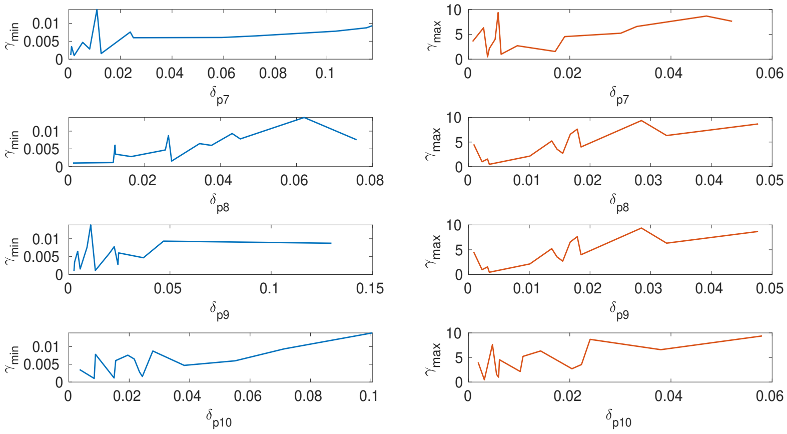


Figure 7: $x$–axis depicts the relative distance of the particular parameter $\theta_i$ from its reference value $\theta_i^*$. $y$–axis on the left is the minimum sum-square deviation and on the right it is the maximum sum-square deviation from $y^*(t)$. The parameters $p_7$ is insensitive for $\delta_i > 0.02$ and $p_9$ for $\delta_i > 0.05$

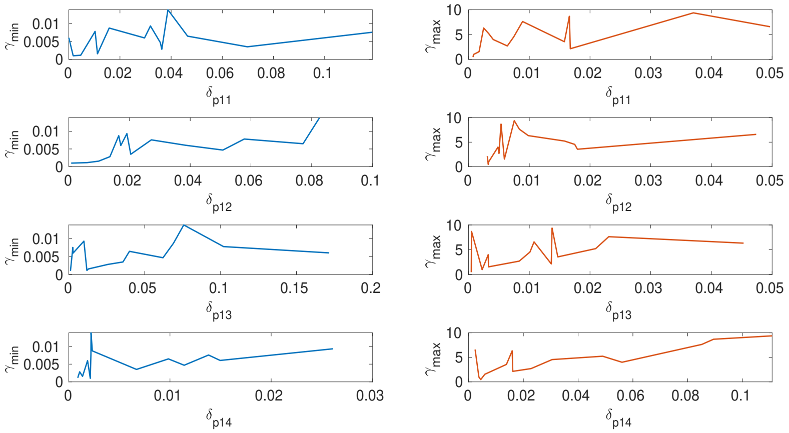


Figure 8: $x$-axis depicts the relative distance of the particular parameter $\theta_i$ from its reference value $\theta_i^*$. $y$-axis on the left is the minimum sum-square deviation and on the it is the right maximum sum-square deviation from $y^*(t)$. The parameters $p_{13}$ and $p_{14}$ are highly sensitive in the vicinity of $\theta_i^*$.

.
