## Supporting information 2 for "Sloppiness: fundamental study, new formalism and its application in model assessment"

This section presents, the necessary model equations used for demonstration of the proposed method.

#### 1 Pharmacodynamic HIV infection mode

The system is defined by four state variables and eight parameters as given below, where $V$ stands for free virus (Model Output).

$$\mathcal{M}:\begin{cases}\dot{T_1} = s - \mu_T T + rT(1 - \frac{(T+T^*+T^{**})}{T_{max}}) - k_1 VT \\ \dot{T^*} = k_1 VT - \mu_T T^* - k_2 T^* \\ \dot{T^{**}} = k_2 T^* - \mu_b T^{**} \\ \dot{V} = N_v \mu_b T^{**} - k_1 VT - \mu_v T\end{cases} \tag{1}$$

#### 2 Mitotic Oscillator

The dynamics of the Mitotic oscillator is governed by the following system differential equations. The system has three states and ten parameters.In the below equations, C denotes the cyclin concentration, while M and X represent the fraction of active cdc2 kinase and the fraction of active cyclin protease.

$$\mathcal{M}:\begin{cases}\frac{dC}{dt} = v_i - k_d C - v_d X \frac{C}{K_d + C} \\ \frac{dM}{dt} = \frac{V1(1-M)}{(K1+(1-M))} - \frac{V_2 M}{K_2 + M} \\ \frac{dX}{dt} = \frac{V3(1-X)}{(K3+(1-X))} - \frac{V_4 X}{X + K_4} \\ y(t) = C(t) + M(t) + X(t) \\ X(0) = M(0) = C(0) = 0.01\end{cases} \tag{2}$$

### 3 Codes

The MATLAB code for constructing visual tool is provided in the following link
`https://github.com/PremJ1704/My_Research/`
